# Coordinated growth of linked epithelia is mediated by the Hippo pathway

**DOI:** 10.1101/2023.02.26.530099

**Authors:** Sophia Friesen, Iswar K. Hariharan

## Abstract

An epithelium in a living organism seldom develops in isolation. Rather, most epithelia are tethered to other epithelial or non-epithelial tissues, necessitating growth coordination between layers. We investigated how two tethered epithelial layers of the *Drosophila* larval wing imaginal disc, the disc proper (DP) and the peripodial epithelium (PE), coordinate their growth. DP growth is driven by the morphogens Hedgehog (Hh) and Dpp, but regulation of PE growth is poorly understood. We find that the PE adapts to changes in growth rates of the DP, but not vice versa, suggesting a “leader and follower” mechanism. Moreover, PE growth can occur by cell shape changes, even when proliferation is inhibited. While Hh and Dpp pattern gene expression in both layers, growth of the DP is exquisitely sensitive to Dpp levels, while growth of the PE is not; the PE can achieve an appropriate size even when Dpp signaling is inhibited. Instead, both the growth of the PE and its accompanying cell shape changes require the activity of two components of the mechanosensitive Hippo pathway, the DNA-binding protein Scalloped (Sd) and its co-activator (Yki), which could allow the PE to sense and respond to forces generated by DP growth. Thus, an increased reliance on mechanically-dependent growth mediated by the Hippo pathway, at the expense of morphogen-dependent growth, enables the PE to evade layer-intrinsic growth control mechanisms and coordinate its growth with the DP. This provides a potential paradigm for growth coordination between different components of a developing organ.

## Results and Discussion

How quickly an epithelium grows, relative to a tissue to which it is tethered, is a fundamental driver of tissue shape. Two tethered epithelia growing at the same rate produce a flat sheet, as in the simple organism *Trichoplax* (Syed & Schierwater, 2002). Differential epithelial growth can cause tissue curvature, as occurs in the vertebrate retina (Moreno-Mármol et al., 2021), and when an epithelium dramatically outgrows neighboring tissues, it can result in buckles and folds, as occurs in the folds of the brain and in the villi of the intestine (Shyer et al., 2013; Tallinen et al., 2014, 2016). Therefore, how tethered epithelia coordinate their growth – or not – is an important developmental problem. Here, we investigate the coordinated growth that occurs between the two epithelial layers of the *Drosophila* wing imaginal disc, the disc proper (DP) and the peripodial epithelium (PE).

The *Drosophila* wing disc is a flattened sac of simple epithelium (Tripathi & Irvine, 2022) composed of two layers that are connected at the edges, similar to a pita pocket. The two layers have distinct morphologies and gene expression patterns throughout the third larval instar (L3) (Auerbach, 1936; White & Wilcox, 1985; Agnes et al., 1999; Pallavi & Shashidhara, 2003, 2005; Nusinow et al., 2008); the DP is made of tightly-packed columnar cells, while the PE consists of a central region of broad, flat squamous cells and a margin of cuboidal cells **(Figure 1A)** (McClure & Schubiger, 2005; Pallavi & Shashidhara, 2005; Tripura et al., 2011). We found that, as the disc grows during L3, the area of the PE, measured by expression of the PE marker gene *eyes absent* (*eya*), stays proportional to the area of the disc as a whole **(Figure 1B-D)**, suggesting that growth of this portion of the PE is coordinated with the rest of the disc.

**Figure 1.**
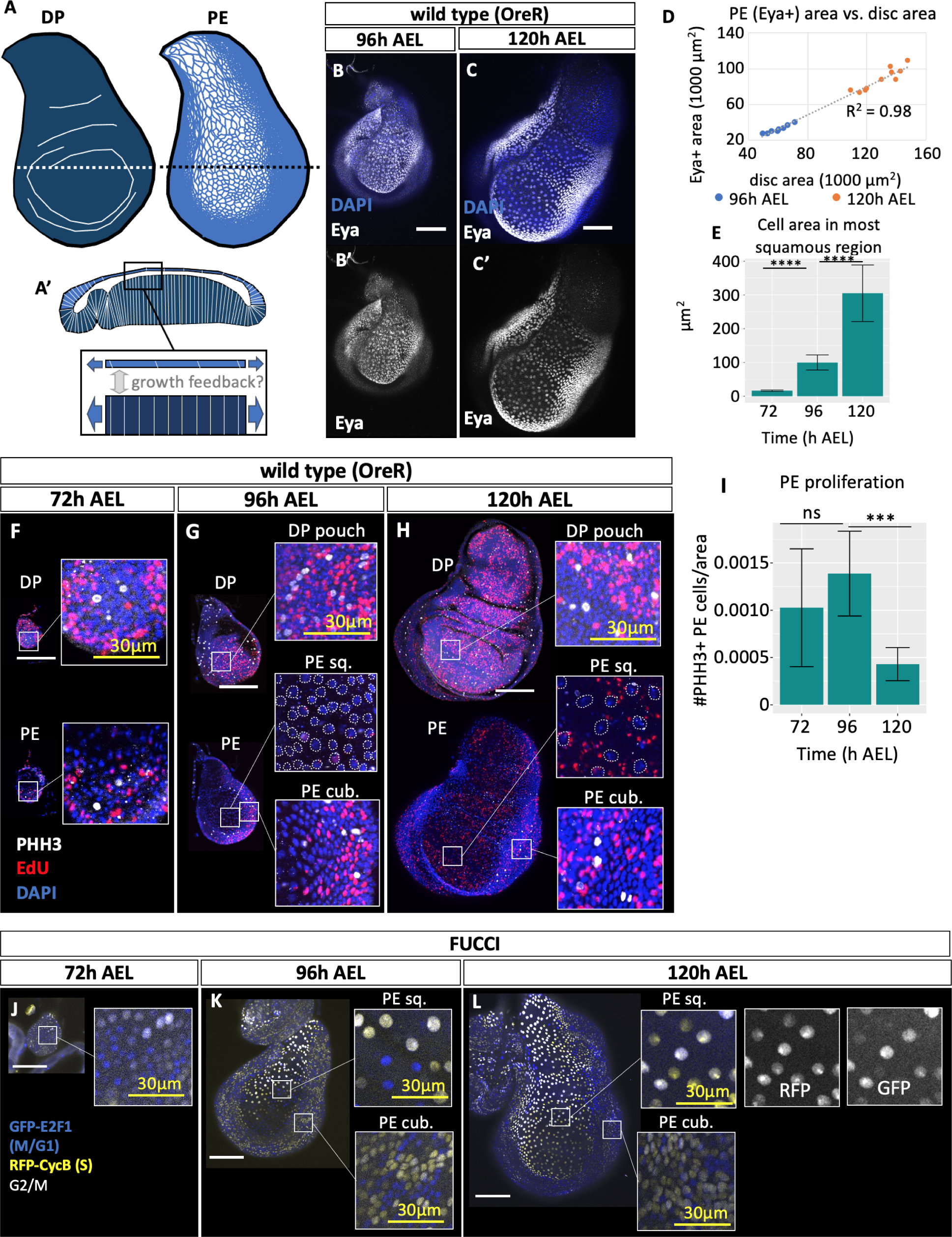
Proportional growth of the PE involves cell shape changes. **A)** Diagram of the two layers of the wing disc in the third larval instar. The disc proper (DP, dark blue) is mostly composed of columnar cells and has characteristic buckles and folds (white lines); the peripodial epithelium (PE, light blue) is composed of squamous cells in the central region and cuboidal cells in more peripheral regions. Representative cell outlines of squamous cells are indicated. **A’)** Diagram showing a cross section through the wing disc along the dotted line in A). Note the differences in cell shape between the two layers. Growth of the two layers could occur independently, or could be regulated by unidirectional or bidirectional communication between the layers. **B)** Mid-L3 (96 h AEL) wild-type wing disc stained for the PE marker Eyes absent (Eya, white). Image is within the plane of the PE. **C)** Late-L3 (120 h AEL) wild-type wing disc stained for Eya, showing that the Eya-positive region remains proportional to the overall size of the disc as the tissue grows. Image is within the plane of the PE. **D)** Quantification of the area of the PE, as approximated by the area of Eya-expressing cells, compared to the total area of the disc, for mid-L3 (96 h after egg lay [AEL]) and late-L3 (120 h AEL) wild-type wing discs. The linear correlation of PE area to total disc area indicates that the PE maintains a constant proportionality to the disc as a whole as the two layers grow. **E)** From early (72 h AEL) to late (120 h AEL) L3, mean cross-sectional area of individual PE cells increases dramatically. Cell area was measured by counting the number of nuclei within the most squamous 50 μm x 50 μm square of the PE and calculating the average area for those cells. n = 11 discs (72h), 14 discs (96h), 13 discs (120h). Unpaired t-test of average cell area between 72h and 96h, ****p<0.00001; unpaired t-test of average cell area between 96h and 120h, ****p<0.00001. **F-H)** The PE proliferates less than the DP, especially in central squamous cells and especially in late L3. Cells in S phase are labeled by EdU incorporation (red), and cells undergoing mitosis are labeled with anti-phospho-histone H3 (PHH3, white). For each time point, the top image shown is in the plane of the DP and the lower image is in the plane of the PE. The DP remains highly proliferative throughout L3. The cuboidal cells at the margins of the PE (PE cub.) continue to proliferate throughout L3, but the squamous cells (PE sq.) stop proliferating, as seen by the absence of EdU and PHH3. Red EdU staining in the squamous PE inset at 120h is fluorescence “bleed-through” from the DP, and does not overlap with PE nuclei (outlined in white in 96h and 120h PE sq. insets). Overall proliferation across the entire PE, as measured by the density of PHH3-positive cells, decreases by late L3. The density of PHH3-positive cells was calculated by counting PHH3-positive cells and dividing by the area of the disc. n = 19 discs (72h), 14 discs (96h), 7 discs (120 h). Unpaired t-test of PHH3-positive cell density between 72h and 96h, ns = not significant (p = 0.061); unpaired t-test of PHH3-positive cell density between 96h and 120h ***p<0.001. **J-L)** Fly-FUCCI shows that the PE progressively arrests in G2. Cells in S phase express RFP-CycB (yellow), cells in late mitosis and G1 express GFP-E2F1 (blue), and cells in G2 and early mitosis express both fluorescent markers (white). **J)** In early L3, PE cells show a “salt-and-pepper” pattern of cell cycle states, indicating asynchronous proliferation. **K)** By mid L3, cuboidal margin cells and some squamous cells show a range of cell cycle states, but a contiguous region of squamous PE cells appears to be synchronized in G2 (white nuclei). **L)** By late L3, most of the squamous PE seems to be arrested in G2, although the cuboidal margin retains a variety of cell-cycle states. All scale bars are 100 μm unless otherwise indicated.

While DP growth in L3 is driven predominantly by cell proliferation, we found that growth of the PE is driven mostly by cell flattening that results in a dramatic increase in cell cross-sectional area **(Figure 1E)** (McClure & Schubiger, 2005). Using EdU incorporation to mark S-phases and anti-phospho-histone H3 (PHH3) to mark mitotic cells, we saw little proliferation within the PE as compared to the DP, especially in the squamous region. While the cuboidal cells at the PE margins continued to proliferate throughout L3, the central squamous region became less proliferative over time **(Figure 1F-I).** Using Fly-FUCCI (Zielke et al., 2014), we found that an increasing number of PE cells in the central squamous region arrest in G2 over the course of L3 **(Figure 1J-L)**. Thus, cell shape changes play a key role in driving the growth of the PE.

Synchronized growth of the PE with the DP could occur through independent growth of the two layers at the same rate, a “leader and follower” model in which growth of one layer sets the pace for the other, or through bidirectional crosstalk in which growth of each layer influences the growth of the other. To distinguish between these models, we experimentally changed the growth rate of one layer and asked whether the other layer remained proportional.

We first manipulated growth in the DP and looked for effects on PE growth **(Figure 2A-F)**. When we reduced DP growth by overexpressing the TOR inhibitors *Tsc1* and *Tsc2* (Tapon et al., 2001; Potter et al., 2001; Gao et al., 2002) within a large region of the DP, PE area was also reduced **(Figure 2A, B, E, F)**. Conversely, overgrowing the DP by expressing the pro-growth microRNA *bantam* (*ban*) (Brennecke et al., 2003) caused PE area to increase proportionally **(Figure 2A, C, E, F).** This indicates that the PE adapts its growth based on growth of the DP. However, when the DP was overgrown by expressing constitutively active Yki (Dong et al., 2007; Oh & Irvine, 2009), the PE did not overgrow to match **(Figure 2A, D, E, F)**, possibly because the rapid overgrowth outpaced the ability of the PE to adjust its growth.

**Figure 2.**
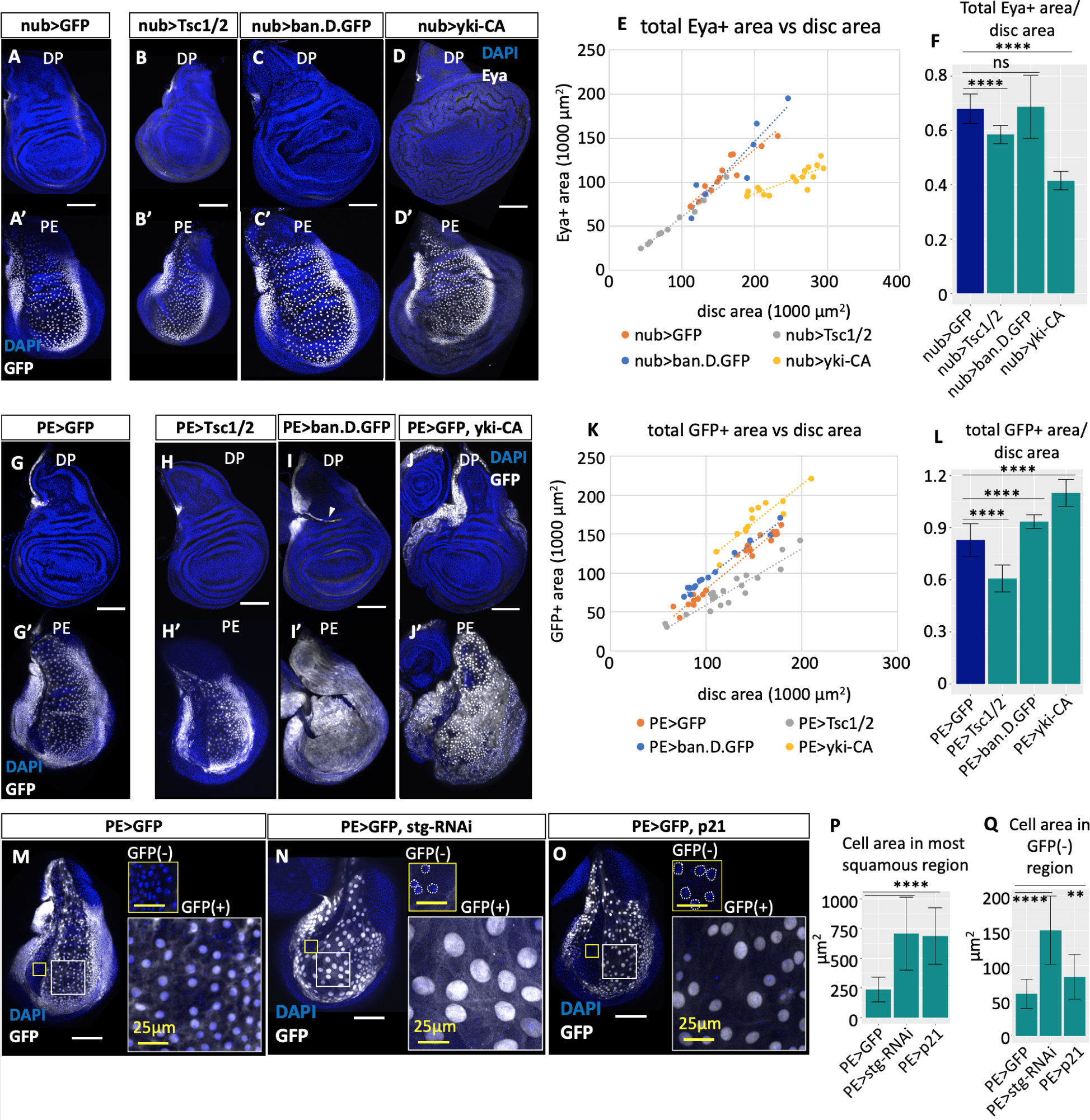
PE growth adapts to the size of the DP and can do so via cell shape changes. **(A-F)** Effect of changing growth of part of the DP on PE growth. All discs are from wandering L3 larvae; all scale bars are 100 μm. The area of the PE is approximated by expression of Eyes absent (Eya, white). **A)** Control disc in which *nubbin-Gal4 (nub-Gal4)* drives *UAS-GFP*. **B)** DP undergrowth caused by *nubbin-Gal4* (*nub-Gal4*) driving *UAS-Tsc1/Tsc2* expression. **C)** Disc in which the DP overgrows due to *nub-Gal4* driving *UAS-bantam.D.GFP*. **D)** Disc in which the DP overgrows severely due to *nub-Gal4* driving constitutively active Yorkie *(UAS-yki-CA)*. **E)** Quantification of PE area as compared to total disc area for the genotypes shown in (A-D). The area of the PE, as approximated by the area of Eya+ tissue, remains roughly proportional to the area of the disc as a whole for most manipulations of DP size; note that most points fall along the same line. However, severe DP overgrowth induced by *nub>yki-CA* does not result in proportional PE overgrowth (yellow points). **F)** The ratio of the area of the PE to the total disc area shows that PE growth adapts to some, but not all, DP growth disruptions. When the DP is overgrown with *ban.D.GFP*, PE size increases proportionally; when the DP is undergrown by overexpression of Tsc1/2, PE size decreases even more than might be expected, despite the lack of any direct genetic manipulation within the PE. However, the PE does not grow proportionally with the DP when the DP is severely overgrown due to *nub>yki-CA.* n = 13 discs *(nub>GFP)*, 10 discs *(nub>Tsc1/2)*, 8 discs *(nub>ban.D.GFP)*, 17 discs *(nub>yki-CA)*. Eya+ area was calculated as the sum of Eya+ area in the plane of the PE and in the plane of the DP. Unpaired t-test of the ratio of Eya+ area to total disc area between *nub>Tsc1/2* and *nub>GFP* ****p<0.0001; unpaired t-test of the ratio of Eya+ area to total disc area between *nub>ban.D.GFP* and *nub>GFP* ns (not significant, p = 0.89); unpaired t-test of the ratio of Eya+ area to total disc area between *nub>yki-CA* and *nub>GFP* ****p<0.00001. **(G-L)** Effect of changing PE growth on growth of the rest of the disc. The area of the PE is approximated by the area of GFP+ expression (white) due to *R15C09-Gal4(PE)>GFP*. **G)** Control disc in which GFP is expressed in the PE using the *PE-Gal4* driver. **H)** When the PE is undergrown due to *PE>Tsc1/2,* the disc proper does not undergrow in response. **I)** Moderate PE overgrowth caused by overexpression of bantam.D.GFP using *PE-Gal4*. Arrowhead indicates where a fold in the disc caused part of the PE to be visible in the plane of the DP. Increased GFP+ area at the anterior margin of the DP suggests part of the PE has “rolled over” into the plane of the DP. **J)** Severe PE overgrowth caused by overexpression of constitutively active Yki (Yki.S168A.V5). Note increased rollover of the PE into the plane of the DP. **K)** Quantification of PE area as compared to total disc area for the genotypes shown in (G-J). The area of the PE, as approximated by the total area of GFP expression, becomes disproportionately small under conditions of PE undergrowth, and disproportionately large under conditions of PE overgrowth, compared to the total disc area. **L)** The ratio of the area of the PE to the total disc area decreases when the PE is undergrown and increases when the PE is overgrown, indicating that the DP does not adjust its growth to remain proportional to the PE. n = 24 (*PE>GFP)*, 19 (*PE>Tsc1/2)*, 15 *(PE>ban.D.GFP)*, 13 *(PE>yki-CA)*. GFP+ area was calculated as the sum of GFP+ area within the plane of the PE and the plane of the DP. Unpaired t-test of the ratio of GFP+ area to disc area between *PE>Tsc1/2* and *PE>GFP* ****p<0.00001; unpaired t-test of the ratio of GFP+ area to disc area between *PE>ban.D.GFP* and *PE>GFP* ****p<0.0001; unpaired t-test of the ratio of GFP+ area to disc area between *PE>yki-CA* and *PE>GFP* ****p<0.00001. **(M-Q)**Cell shape changes can drive adaptive PE growth. **M)** Control disc in which GFP (white) is expressed on the *PE-Gal4* driver. White scale bars are 100µm; yellow scale bars in insets are 25µm. **N)** When cell division within the PE is inhibited by knocking down *string* on the *PE-Gal4* driver, the PE contains many fewer cells, but maintains proportionality with the DP via a dramatic increase in cell area. Nuclei in GFP(-) inset are outlined in white. The increased area of GFP+ cells is likely the result of endoreplication caused by *stg-RNAi*. **O)** Inhibiting the G1-S transition by overexpressing p21 on the *PE-Gal4* driver similarly reduces the number of cells in the PE, but PE area adapts via an increase in cell area. Nuclei in GFP-inset are outlined in white. **P)** Quantification of cell area for genotypes shown in **(M-O)**. Average cell area of squamous cells was calculated by dividing the area of the most squamous 2500μm square region of the PE by the number of nuclei in that region. n = 10 (*PE>GFP*), 16 (*PE>stg-RNAi*), 12 (*PE>p21*). Unpaired t-test of average cell area between *PE>GFP* and *PE>stg-RNAi* ****p<0.0001; unpaired t-test of average cell area between *PE>GFP* and *PE>p21* ****p<0.0001. **Q)** Quantification of cell area within the GFP-negative region of the PE for genotypes shown in **(M-O)**. n = 15 *(PE>GFP)*, 12 *(PE>stg-RNAi),* 22 *(PE>p21).* Unpaired t-test of average cell area between *PE>GFP* and *PE>stg-RNAi* ****p<0.00001; unpaired t-test of average cell area between *PE>GFP* and *PE>p21* **p<0.01.

To determine whether the DP adjusts its growth in response to growth of the PE, we altered growth specifically within the PE **(Figure 2G-L)**. We found that the two drivers most commonly used to manipulate gene expression in the PE, *Ubx-Gal4* (Pallavi & Shashidhara, 2003) and *AGiR-Gal4* (Gibson et al., 2002), both had significant expression in the DP **(supplemental fig. S1A, B)**. We therefore used the previously uncharacterized FlyLight driver *R15C09-Gal4* (Jenett et al., 2012) (henceforth referred to as *PE-Gal4*), which is expressed in the majority of the PE throughout L3 and which has minimal DP expression **(supplemental fig. S1C-E)**.

When we reduced PE growth by overexpressing both *Tsc1* and *Tsc2*, the DP did not undergrow to match **(Figure 2G, H, K, L)**. Conversely, *ban* expression within the PE caused the area of the PE, as measured by driver expression, to increase disproportionately to the size of the disc as a whole **(Figure 2G, I, K, L)**. In contrast to the results of overexpressing *ban* or *Tsc1* and *Tsc2* in the DP, this disproportionality shows that the DP does not adjust its growth to match changes in PE growth. More severe overgrowth, driven by expression of constitutively active Yki **(Figure 2G, J, K, L)**, brought the PE even further out of proportion, as more PE cells “rolled over” into the plane of the DP. Thus, at least within a certain range of growth rates, growth of the DP seems to set the pace for growth of the PE, but not vice versa.

Since cell shape changes are a major contributor to normal PE growth, we wondered whether cell shape changes alone could allow adaptive PE growth. To explore this possibility, we knocked down the Cdk1 activator *string (stg)* in the PE, which is expected to block mitosis (Edgar & O’Farrell, 1990; Mozer & Easwarachandran, 1999). While this greatly reduced the number of PE cells, the area of the PE remained largely proportional to the area of the disc as a whole, due to a nearly three-fold increase in cell area **(Figure 2M, N, P, Q)**. Similarly, overexpression of the human Cdk inhibitor *p21*, which inhibits the G1/S transition in *Drosophila* (de Nooij & Hariharan, 1995), reduced PE cell number but increased PE cell area, such that the PE was only slightly smaller than normal **(Figure 2M, O, P, Q)**. Therefore, adaptive growth that occurs mostly by an increase in cell area can allow the PE to match DP growth.

Importantly, reducing the number of cells in the PE increased cell area not only in the cells directly impacted by cell cycle inhibition, but also increased the area of adjacent PE cells within an area at the AP boundary where *PE-Gal4* is not expressed (GFP(-) insets in **Figure 2M-O**, quantified in **Figure 2Q**). The increased area of PE cells near to, but outside of, the *PE-Gal4* domain indicates that cell shape changes are not directly caused by cell-autonomous cell-cycle perturbations, but rather by a separate signal, such as increased mechanical stretch, that extends beyond *Gal4-*expressing cells.

Our results show that the PE adjusts its growth to match the DP, but what regulates growth of the PE? Key to DP growth are two morphogens, Hedgehog (Hh) and Decapentaplegic (Dpp) (Tabata & Kornberg, 1994; Burke & Basler, 1996; Strigini & Cohen, 1997; Affolter & Basler, 2007; Restrepo et al., 2014). In the DP, Hh is expressed throughout the posterior compartment and moves into the anterior compartment to activate expression of its target *dpp* along the compartment boundary **(Figure 3A)** (Zecca et al., 1995). Dpp spreads across the DP to promote cell survival and proliferation (Martín-Castellanos & Edgar, 2002; Rogulja & Irvine, 2005); clones of cells in the DP that cannot receive Dpp undergrow or die outright (Burke & Basler, 1996; Gibson & Perrimon, 2005; Wartlick et al., 2011). Both Hh and Dpp are expressed in the PE in a similar pattern to that found in the DP **(Figure 3A, supplemental fig. S2A, B, L)** (Pallavi & Shashidhara, 2005; Tang et al., 2016), although Hh expression in the PE is faint enough to have escaped detection with some methods (Pallavi & Shashidhara, 2005).

**Figure 3.**
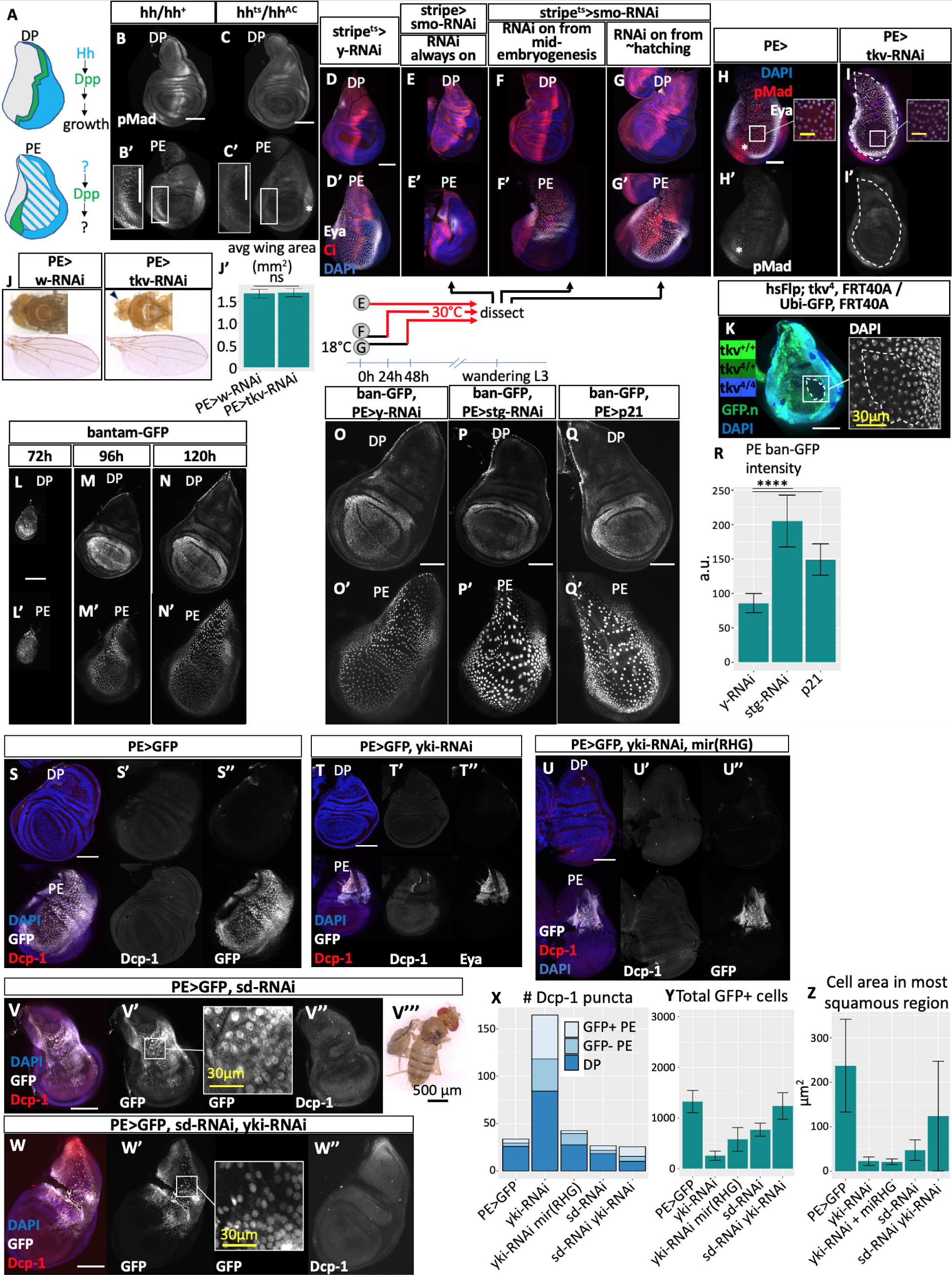
Mechanically-driven growth is more important than morphogen-driven growth in the PE. **A)** Diagram of Hh and Dpp expression in the DP and PE. Hh (light blue) is expressed in the posterior compartment of each layer, although expression in the squamous PE is faint and difficult to detect (blue stripes). Dpp (green) is expressed in a narrow stripe in both the DP and the PE. Note the anterior shift of the compartment boundary in the PE. **B)** Dpp signaling at the PE compartment boundary requires Hh. Dpp signaling is visualized by staining for phospho-Mad (white). In *hh/+* discs, Dpp signaling is visible at the AP boundary of the DP (top) and the PE (bottom and inset). **C)** Discs without functional Hh, which have a *hh* null allele (*hh^AC^*) and a temperature-sensitive *hh* allele (*hh^ts^*), lose Dpp signaling at the AP boundary in both layers after 15 h at the nonpermissive temperature (30°C). Asterisk in **C’** denotes a posterior patch of Hh-independent Dpp activity as previously described(Foronda et al., 2009). (**D-G**) Hh signaling is required for very early PE growth but not later growth. PE cells are labeled by Eya staining (white); anterior compartment is labeled by Ci staining (red). **D)** Control disc expressing *yellow-RNAi* on the *stripe-Gal4* driver. Note anterior Ci expression and the large Eya+ region. **E)** Constitutive knockdown of *smo* at the AP boundary of the PE causes severe PE undergrowth. **F)** When *smo* knockdown is initiated at mid-embryogenesis, after 24h at 18°C, mild PE undergrowth occurs. **G)** *smo* knockdown beginning around larval hatching at 48h at 18°C does not cause PE undergrowth. (**H-I**) Dpp signaling is dispensable for PE growth. **H)** Control disc showing Dpp signaling as observed by pMad staining (red in H, white in H’.) White scale bar is 100µm; yellow scale bar is 30µm. **I)** Knockdown of the Dpp receptor *tkv* in the PE abolishes detectable Dpp signaling. Dotted line indicates knockdown domain. Asterisk indicates region of high pMad in control discs. The *PE>tkv-RNAi* disc PE grows to large size as observed by Eya expression (white in top panels), and cells are appropriately squamous as assessed by spacing of nuclei. White scale bar is 100µm; yellow scale bar in inset is 30µm. **J)** Loss of Dpp signaling in the PE does not interfere with its functions of thorax closure and wing eversion during metamorphosis. *PE>tkv-RNAi* adults have normal dorsal thoraces and wings. Note that *PE-Gal4* also expresses in the eye disc and that *tkv*-*RNAi* leads to substantial undergrowth in this context (arrowhead), confirming knockdown efficacy. **(J’)** Wing area (averaged between left and right wings of female flies) is unchanged by *PE>tkv-RNAi.* n = 7 flies (*PE>w-RNAi*), 12 flies (*PE>tkv-RNAi*). Unpaired t-test of wing area between *PE>w-RNAi* and *PE>tkv-RNAi* p = 0.66 (not significant, ns). **K)** Clones of PE cells that cannot receive Dpp signaling can still proliferate and become squamous. In a disc heterozygous for the loss of function *tkv* allele *tkv^4^*, clones lacking functional *tkv* contain no GFP (no green) and wild-type twinspot clones contain two copies of GFP (bright green). White dotted line indicates the border of a large *tkv^4/4^*clone. *tkv^4/4^* clones in the PE can grow to large size and become squamous (note widely spaced nuclei labeled by DAPI, white in inset). White scale bar is 100µm; yellow scale bar is 30µm. **L)** Yki activity, as measured by expression of the *bantam-GFP* reporter, is high in the PE throughout L3. Note bright squamous nuclei visible at 72h AEL. **M)** Yki activity is high in the PE at 96h AEL. **N)** Yki activity is high in the PE at 120h AEL. **(O-R)** Inhibiting the cell cycle in the PE causes not only increased cell area, but an accompanying increase in Yki activity as measured by bantam-GFP (white). **O)** Control disc expressing bantam-GFP and *PE>y-RNAi*. **P)** When mitosis is inhibited in the PE using *PE>stg-IR, ban-GFP* expression in the PE increases. **Q)** When DNA synthesis is inhibited in the PE using *PE>p21*, *ban-GFP* expression in the PE increases. **R)** Average fluorescence intensity of ban-GFPexpressing nuclei across the area of the PE for control (*PE>y-RNAi)* discs and for discs in which the cell cycle is partly inhibited *(PE>stg-RNAi* and *PE>p21)*, as shown in (O-Q). n = 7 discs (*PE>y-RNAi)*, 18 discs *(PE>stg-RNAi),* 14 discs *(PE>p21)*. Unpaired t-test of PE GFP intensity between *PE>y-RNAi* and *PE>stg-RNAi* ****p<0.00001; unpaired t-test of PE GFP intensity between *PE>y-RNAi* and *PE>p21* ****p<0.00001. **S)** *PE>GFP* control disc showing expression of GFP and the apoptosis marker Dcp-1 (red in **S**, white in **S’**). Note that Dcp-1 staining is essentially absent from wild-type PE. **T)** Knockdown of *yki* in the PE causes severe PE undergrowth as observed by an extremely reduced area of Eya expression (white), as well as increased apoptosis in both layers of the disc (Dcp-1, red in **T**, white in **T’**). Yki-deficient PE also lacks squamous cells. **U)** *yki* knockdown reduces PE cell number and prevents PE cell shape changes, even in the absence of cell death. Apoptosis in the PE was blocked by expression of the microRNA against reaper, hid, and grim (*mir(RHG)*), concurrent with Yki knockdown. Despite a lack of apoptosis in the PE (Dcp-1, red in **U**, white in **U’**), the Yki-deficient PE (marked by GFP, white) contains fewer cells and is not squamous. **V)** Knockdown of the Yki cofactor scalloped (*sd*) in the PE causes reduced cell number and reduced cell area, but does not cause increased apoptosis. Note the smaller area of GFP+ tissue (white in left panels and inset) in the *sd* knockdown condition (compare to **2G**) and increased density of nuclei corresponding to reduced cell area. Apoptosis, as measured by staining for Dcp-1 (red in left panel, white in right panel), was not increased by *sd* knockdown. *PE>sd-RNAi* causes near-complete lethality (6 adults eclosed out of 233 pupae), and rare escapers show wing eversion defects. **W)** Simultaneous knockdown of both *yki* and *sd* in the PE causes a *sd-RNAi-*like phenotype, with reduced GFP+ tissue area (white in left panels and inset) but no increased apoptosis as measured by Dcp-1 (red in left panel, white in right panel), indicating that all of Yki’s growth effects in the PE are dependent on Sd. **X)** Quantification of cell death in experiments shown in panels **(S-W)**, showing increased apoptosis upon Yki knockdown and successful block of apoptosis upon coexpression of mir(RHG). n = 9 discs (TM2), 7 discs *(yki-RNAi)*, 5 discs *(yki-RNAi mir(RHG))*, 10 discs *(sd-RNAi)*, 8 discs *(sd-RNAi yki-RNAi)*. **Y)** Quantification of cell number in *yki-RNAi* PE compared to control discs and discs coexpressing *mir(RHG)*. Expression of UAS-GFP on *PE-Gal4* is used as a proxy for PE identity. *yki-RNAi* and *yki-RNAi, mir(RHG)* PE contain fewer cells than control PE, indicating that Yki is required for PE proliferation as well as cell survival. n = 6 discs (TM2), 9 discs (*yki-RNAi)*, 7 discs (*yki-RNAi mir(RHG)*), 8 discs *(sd-RNAi)*, 8 discs (*sd-RNAi yki-RNAi*). **Z)** Quantification of cell area in *yki-RNAi* PE compared to control discs and discs coexpressing *mir(RHG)*. *yki* knockdown prevents cells from becoming squamous, even in the absence of cell death. Average cell area was calculated by counting nuclei in the most squamous 50×50 square region of each disc and multiplying the reciprocal by 2500um^2^. n = 10 discs (TM2), 15 discs (*yki-RNAi*), 7 discs (*yki-RNAi mir(RHG)*), 11 discs (*sd-RNAi*), 8 discs (*sd-RNAi yki-RNAi*). All scale bars are 100µm unless otherwise stated.

If Hh and Dpp function as layer-intrinsic regulators of growth, as their expression patterns suggest, then how can the PE adjust its growth to match the DP? We investigated whether Hh and Dpp regulate PE growth.

To test whether Dpp expression in the PE requires *hh,* we inactivated *hh* throughout 3^rd^-instar larvae using the temperature-sensitive *hh^ts2^* allele (Ma et al., 1993). In wing discs without functional *hh*, we observed a loss of Dpp signaling as assessed by phosphorylated Mad (pMad) levels (Wiersdorff et al., 1996; Wisotzkey et al., 1998) at the compartment boundary in both tissue layers, showing that Dpp signaling in the PE is Hh-dependent **(Figure 3B, C, supplemental fig. S2F, G)**. Global Hh loss also reduced proliferation in both layers **(supplemental fig. S2C, D, E)**. Organism-wide Hh loss has been reported to affect PE cell shape (McClure & Schubiger, 2005), but this would be expected given the requirement for Hh in the DP and given the coordination of PE growth with the DP.

We next asked whether Hh signaling is required in the PE itself by knocking down the Hh pathway component *smoothened* (*smo)* in a stripe along the PE compartment boundary, where we expected Hh signaling to be strongest (Basler & Struhl, 1994; Chen & Struhl, 1996). We used a different PE-specific driver, *R43F06-Gal4* (Jenett et al., 2012), henceforth referred to as *stripe-Gal4*, which expresses along the AP compartment boundary of the PE throughout L3 **(supplemental fig. S1F, G, H, S2H)**. Expression of *stripe>smo-RNAi* throughout development caused severe PE undergrowth, indicating that Hh is indeed required for PE growth at some point in development **(Figure 3D, E, supplemental fig. S2K)**. However, when we used a temperature-sensitive Gal80 to prevent RNAi expression before mid-embryogenesis, PE undergrowth was much less severe **(Figure 3D, F, supplemental fig. S2K)**, and when RNAi expression began at larval hatching, the PE was of normal size **(Figure 3D, G, supplemental fig. S2J, K)**. In contrast, similarly timed *smo* knockdown using a DP-specific driver caused significant DP undergrowth **(supplemental fig. S2I)**. This shows that Hh signaling is necessary for PE growth or development during embryogenesis. However, unlike in the DP, Hh signaling at the compartment boundary is dispensable for PE growth throughout the larval stages.

Inhibition of Dpp activity within the PE has been reported to prevent squamous cell morphogenesis (McClure & Schubiger, 2005); however, prior experiments using *Ubx-Gal4* likely impacted Dpp signaling and growth within the DP as well (see **supplemental fig. S1A)**, raising the question of whether Dpp signaling is specifically required in the PE. We investigated whether Dpp signaling is required for PE growth by knocking down the Dpp receptor *thickveins (tkv)* (Brummel et al., 1994; Ruberte et al., 1995; Nellen et al., 1996) throughout most of the PE. *PE>tkv-RNAi* reduced Dpp signaling in the PE to undetectable levels, but the PE became large and squamous regardless **(Figure 3H, I)**, and adult flies eclosed without the wing or thoracic midline defects which, if they occurred, might suggest PE dysfunction (Milner et al., 1984; Agnes et al., 1999; Pastor-Pareja et al., 2004; Tripura et al., 2011; Kosakamoto et al., 2018) **(Figure 3J, J’)**, indicating that the *tkv*-deficient PE functions normally during metamorphosis. To confirm that Dpp signaling is not required for PE growth, we made *tkv* null clones throughout the wing disc using mitotic recombination. *tkv* null clones in the DP were eliminated, and in the PE, these clones usually contained fewer cells than their wild-type twin-spots, implying some growth disadvantage **(supplemental fig. S2N)** (Adachi-Yamada & O’Connor, 2002; Gibson et al., 2002; Moreno et al., 2002). However, such clones were still able to grow to a large size and *tkv* null cells still became squamous **(Figure 3K)**. Therefore, Dpp and Hh signaling are not critical regulators of either PE growth or cell shape changes.

We noticed that the expression of Dpp target genes *Daughters against Dpp (Dad)* (Tsuneizumi et al., 1997) and *brinker (brk)* (Minami et al., 1999; Tang et al., 2016) was patterned in the PE **(supplemental fig. S2L, M)**, prompting us to ask if Dpp regulates patterned gene expression in the PE, as it does in the DP (Nellen et al., 1996; Morimura et al., 1996; Lecuit et al., 1996; Haerry et al., 1998). Overexpression of *dpp* using *PE-Gal4* strongly reduced Eya expression **(supplemental fig. S2O, P)**, while reducing Dpp signaling using *tkv-RNAi* caused ectopic Eya expression **(Figure 3H, I, supplemental fig. S2Q, R)**, showing that Dpp still patterns gene expression in the PE in a dose-dependent manner. Thus, the roles of Dpp in regulating growth and in patterning gene expression have been uncoupled in the PE.

Mechanical stretching promotes cell proliferation in the wing disc (Schluck et al., 2013). Given the apparent unimportance of Dpp in PE growth, we wondered if growth of the DP could induce stretching of the PE, which could be sensed and responded to by a mechanosensitive growth regulatory pathway such as the Hippo pathway (Zhao et al., 2007; Dupont et al., 2011; Rauskolb et al., 2014; Benham-Pyle et al., 2015; Fletcher et al., 2018; Pan et al., 2018; Chang et al., 2020). The Hippo pathway component Yorkie (Yki) is a transcriptional coactivator which promotes growth in many contexts (Huang et al., 2005; Dong et al., 2007; Davis & Tapon, 2019; Zheng & Pan, 2019). Additionally, Yki has been reported to be more active in the PE, and Yki deficiency in the PE results in the absence of squamous cells (Borreguero-Muñoz et al., 2019; Fletcher et al., 2018), although it is unclear from these prior studies whether the lack of squamous cells is the result of cell death, reduced proliferation, or a role for the Hippo pathway in PE cell shape changes. This prompted us to investigate how the Hippo pathway regulates PE growth.

We observed high levels of the Yki activity reporter *bantam-GFP (ban-GFP)* (Matakatsu & Blair, 2012) in the PE throughout L3 **(Figure 3L, M, N)**. If increased Yki activity is caused by PE stretching, then conditions that result in excessive stretching of PE cells should further increase Yki activity. Indeed, *ban-GFP* expression was highly upregulated by *PE>stg-RNAi* or *PE>p21*, both of which inhibit cell proliferation and result in fewer, larger-area PE cells **(Figure 3O, P, Q, R)**. This result indicates that elevated Yki activity is correlated with tissue stretch, consistent with prior findings (Benham-Pyle et al., 2015; Dupont et al., 2011; Rauskolb et al., 2014; Fletcher et al., 2018).

To determine whether PE growth requires *yki*, we knocked down *yki* in the PE. Consistent with published results (Fletcher et al., 2018), *yki* deficiency caused severe PE undergrowth and an absence of squamous cells **(Figure 3S, T**, quantified in **Figure 3Y, Z)**. We observed substantial apoptosis in *PE>yki-RNAi* discs (quantified in **Figure 3X**), which demonstrated that *yki* is required for PE cell survival, and which could mask additional functions for Yki in PE growth.

To determine whether *yki* plays a role in PE development beyond cell survival, we blocked apoptosis in the PE using *mir(RHG)* (Siegrist et al., 2010), which inactivates the three pro-apoptotic genes *reaper*, *hid,* and *grim*, and simultaneously knocked down *yki* **(Figure 3S, U)**. While a much larger region of PE cells survived in such discs, they still contained fewer PE cells than control discs **(Figure 3Y)**, showing that *yki* also promotes PE cell proliferation. Importantly, despite the PE as a whole being disproportionately small, *yki-RNAi mir(RHG)* cells were not squamous **(Figure 3Z)**. This shows that mechanical forces caused by growth discrepancies between the layers are not sufficient to make cells squamous in the absence of *yki*. Instead, our results suggest a specific function for *yki* in making cells squamous in response to stretch.

Yki is a transcriptional co-activator that acts mostly by binding to the sequence-specific DNA-binding protein Scalloped (Sd) and activating gene expression (Wu et al., 2008; L. Zhang et al., 2008; Goulev et al., 2008). In the absence of Yki, Sd binds to repressors and is thought to reduce expression of target genes (Guo et al., 2013; Koontz et al., 2013). Loss of *sd* causes the loss of both its activator and repressor functions and often results in relatively minor changes in tissue size (Goulev et al., 2008; Wu et al., 2008; L. Zhang et al., 2008; Kowalczyk et al., 2022).

Although *sd* null clones fail to grow in the PE of the eye disc (Kowalczyk et al., 2022), and Yki/Sd-mediated cell proliferation promotes stretch-induced growth of the egg follicle epithelium (Borreguero-Muñoz et al., 2019), the frequent lack of a strong growth phenotype upon *sd* loss has called into question whether the Hippo pathway plays a significant role in size regulation during normal development (Kowalczyk et al., 2022).

We therefore asked whether Sd is required for PE growth by knocking *sd* down throughout the PE. *PE>sd-RNAi* caused PE undergrowth and reduced cell number **(Figure 3S, V, Y)**, although *sd* deficiency did not increase apoptosis **(Figure 3X)**. Notably, cells were less squamous when *sd* function was reduced **(Figure 3Z)**. *PE>sd-RNAi* was mostly lethal, and the few (6/233) adults that eclosed had wing eversion defects, indicating that Sd-mediated Hippo signaling is required both for the normal growth of the PE as well as its role in wing eversion. Combined knockdown of both *sd* and *yki* produced mild undergrowth in the PE, similar to *sd* knockdown alone and distinct from the more severe undergrowth caused by *yki* knockdown alone **(Figure 3W, X, Y)**. The *sd-RNAi*-like phenotype of *PE>yki-RNAi sd-RNAi* suggests that all of *yki*’s growth effects in the PE depend on *sd*.

How does the Yki/Sd complex promote growth of the PE? In the DP, an immediate downstream target of Yki/Sd is the microRNA *bantam* (*ban*), which promotes proliferation and inhibits apoptosis (Brennecke et al., 2003; Nolo et al., 2006; Thompson & Cohen, 2006). Moreover, *ban*-*GFP* is highly expressed in the PE **(Figure 3L-N)**. To ask if the growth-promoting function of *yki* in the PE is mediated through *ban*, we generated *ban* null clones. Such clones were rare in the PE **(Figure 4A)**, suggesting that *ban* is required for PE cell survival or growth. Though rare, *ban* null PE cells were appropriately squamous. *ban* overexpression caused modest PE overgrowth, increasing total cell number and the relative size of the affected tissue **(Figure 4B)**. *ban* overexpression was able to rescue most, but not all, of the effects of *yki* knockdown, including total cell number and tissue size **(Figure 4C, O)**. Additionally, *ban-*overexpressing *yki-RNAi* discs had levels of apoptosis comparable to discs overexpressing *ban* alone **(Figure 4M)**. However, *bantam* overexpression did not restore squamous cell shape to *yki-RNAi* discs **(Figure 4C, N)**. Thus, while *bantam* can mediate Yki’s proliferative and anti-apoptotic functions, it is neither necessary nor sufficient to induce the cell shape changes that occur in the PE.

**Figure 4.**
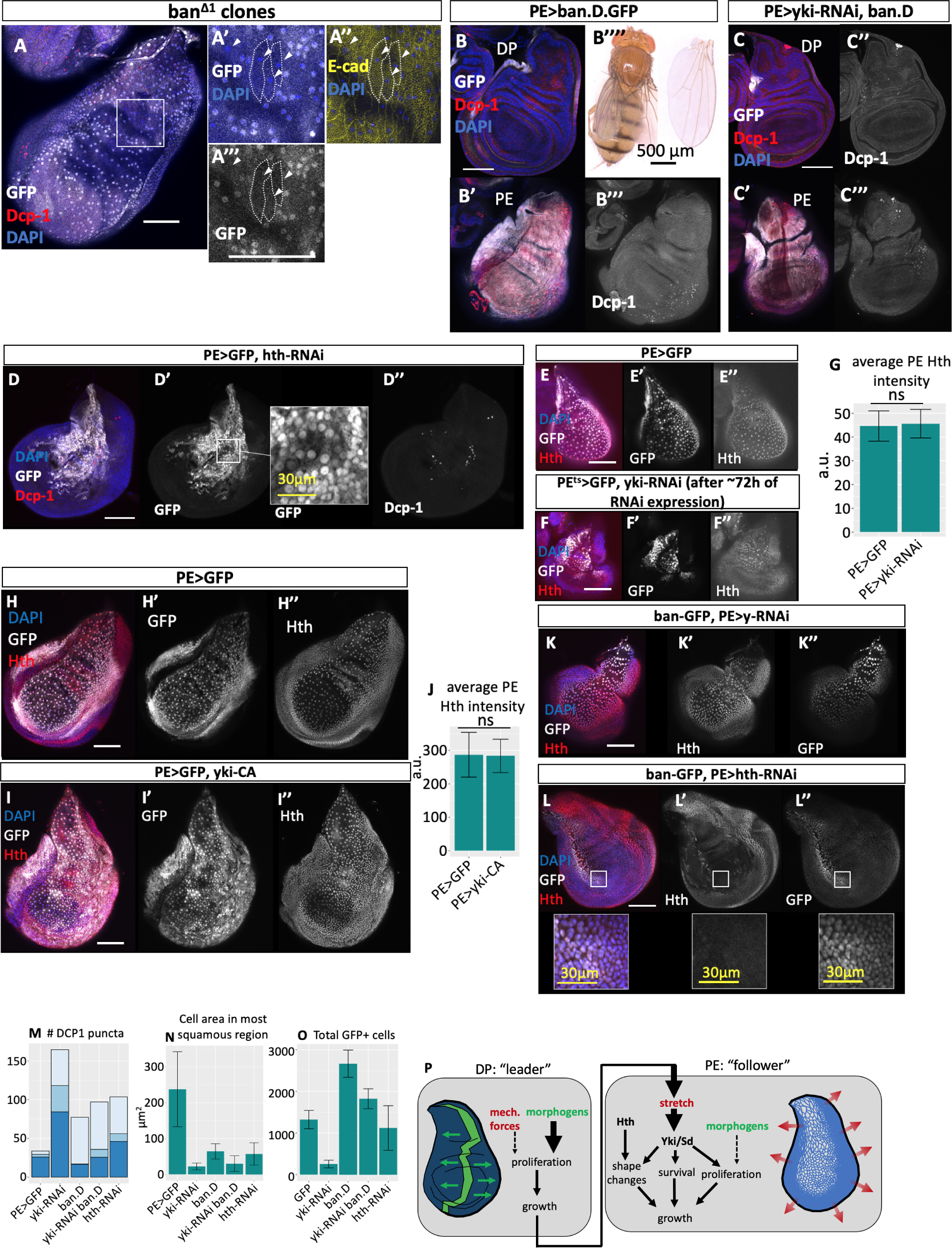
*bantam* and *homothorax* regulate PE growth. **A)** *bantam* null clones (GFP-, arrows in insets) are rarely recovered in the PE, but achieve squamous morphology comparable to their neighbors. Cells are outlined to show their shape (dotted lines) which can be seen by E-cadherin staining (yellow, rightmost panel). **B)** *bantam* overexpression in the PE causes increased cell number (quantified in **4O**) and a corresponding decrease in cell area (quantified in **4N**). *PE>ban-D-GFP* also results in elevated apoptosis in the distal posterior margin of the PE (Dcp-1 staining, red in left panels, white in right panel) (quantified in **4M**). **C)** *bantam* overexpression in the PE largely rescues overall tissue size as well as cell number in *yki-RNAi* discs, and leads to a pattern of apoptosis resembling that caused by *bantam* overexpression alone (quantified in **4M, O**). However, *bantam* overexpression does not rescue squamous cell shape in *yki-RNAi* wing discs (quantified in **4N**). **D)** Knockdown of the putative Yki cofactor *homothorax (hth)* in the PE causes reduced cell area (note inset in **D’**, quantified in **4N**), reduced PE area (as approximated by *PE>GFP,* white in **D** and **D’)**, and increased apoptosis (Dcp-1 staining, red in **D**, white in **D’’,** quantified in **4M-O**). **E)** Hth protein (red in **E,** white in **E’’**) is expressed in the PE. **F)** Hth expression in the PE does not require Yki. When *yki* is knocked down 3 days prior to dissection, Hth protein (red in **F,** white in **F’’**) can still be detected in the PE at comparable levels to wild-type. Knockdown domain is indicated by expression of GFP (white in **F** and **F’**). Discs were imaged and stained in parallel with those in **4E.** **G)** Quantification of Hth expression in the PE for conditions shown in **4E** and **4F**. Average Hth intensity was measured across the area of the disc within the plane of the PE. n= 8 discs (*PE>GFP)*, 6 discs *(PE>yki-RNAi*). Unpaired t-test of Hth intensity between *PE>GFP* (control) and *PE>yki-RNAi* ns (not significant, p = 0.77). **H)** Hth (red in **H,** white in **H’’**) is expressed throughout the wild-type PE. **I)** Ectopic Yki activity is not sufficient to upregulate Hth expression in the PE. When constitutively active Yki is expressed in the PE using *PE-Gal4*, Hth staining intensity in the PE remains unchanged (red in **I**, white in **I’’**). Discs were imaged and stained in parallel with those in **4H**. **J)** Quantification of Hth expression in the PE for conditions shown in **4H** and **4I**. Average Hth intensity was measured across the area of the disc within the plane of the PE. n= 11 discs (*PE>GFP)*, 12 discs *(PE>yki-CA)*. Unpaired t-test of Hth intensity between *PE>GFP* (control) and *PE>yki-CA* ns (not significant, p = 0.89). **K)** Control discs expressing ban-GFP and *PE>y-RNAi.* Note strong expression of both Hth (red in **K,** white in **K’’**) and ban-GFP (white in **K** and **K’’**) throughout the PE. **L)** Yki activity in the PE, as measured by ban-GFP expression, does not require *hth. PE>hth-RNAi* greatly reduces Hth expression (red in **L,** white in **L’**) within the knockdown domain, but ban-GFP expression (white in **L** and **L’’**) remains strong even in regions lacking Hth (see insets). **M)** Quantification of apoptosis in the PE and the DP under various Hippo pathway manipulations (shown in **4B-D** and **3S-T**). Note that *PE>GFP* and *yki-RNAi* data is also shown in **3X**, and is shown here for ease of comparison. Apoptosis was quantified by the number of Dcp-1-positive puncta, counted separately in the DP (dark blue), the GFP-negative PE (where *PE-Gal4* is not expressed, medium blue), and the GFP-positive PE (where *PE-Gal4* is expressed, pale blue). Apoptosis is increased by knockdown of either *yki* or *hth,* or by overexpression of *bantam. bantam* overexpression in *yki-RNAi* causes a pattern of apoptosis most similar to *bantam* overexpression alone. **N)** Quantification of cell area in the squamous PE under various Hippo pathway manipulations (shown in **4B-D** and **3S-T**). Note that *PE>GFP* and *yki-RNAi* data is also shown in **3X**, and is shown here for ease of comparison. Cell area was calculated by dividing the area of the most squamous 50×50μm square region of the PE by the number of cell nuclei in that region. Like *yki-RNAi,* depletion of *hth* causes reduced cell cross-sectional area, though to a lesser extent. *bantam* overexpression also reduces the area per cell, likely due to an increase in cell number, and is unable to rescue squamous cell shape upon *yki* knockdown. **O)** Quantification of total GFP+ cell number (as an approximation of PE cell number) under various Hippo pathway manipulations (shown in **4B-D** and **3S-T**). Note that *PE>GFP* and *yki-RNAi* data is also shown in **3X**, and is shown here again for ease of comparison. *yki-RNAi* reduces PE cell number, an effect that is rescued by coexpression of *bantam. bantam* overexpression increases cell number in the PE. **P)** Proposed model of growth synchronization between the PE and the DP, in which morphogen-driven DP growth mechanically stretches the PE. Stretch forces are sensed and responded to by Hippo signaling via Yki/Sd, which together with Hth, ultimately allows the PE to match its growth to the DP. All scale bars are 100µm unless otherwise stated.

The transcription factor Homothorax (Hth) has been reported to act with or downstream of Yki to regulate *ban* and specify eye disc PE (Peng et al., 2009; T. Zhang et al., 2011). Although the double-knockdown results of *sd-RNAi yki-RNAi* suggest that Sd is the only important Yki cofactor in the wing disc, we asked whether Hth also regulates PE growth. *hth* knockdown in the wing PE caused increased apoptosis and prevented cells from becoming properly squamous **(Figure 4D, M, N)**, although cell proliferation was unaffected **(Figure 4O)**. Hth protein expression levels are independent of Yki activity levels **(Figure 4E-J)**, and *bantam-GFP* expression does not require Hth **(Figure 4K, L)**. These results suggest that, unlike in the eye disc, Hth in the wing disc operates in parallel to Yki/Sd. Thus, we have identified two separate regulators of PE growth, Hth and Yki/Sd, which act independently to promote PE growth via cell shape changes.

Morphogen-driven growth appears to be the default mode of growth within *Drosophila* imaginal discs, and the Hh/Dpp signaling axis appears to exist and retain some patterning functions even within the PE of the wing disc. However, relative to the DP, we find that growth of the PE is regulated primarily by the mechanosensitive Hippo pathway, and is largely independent of morphogen signaling, a difference that allows the PE to match its growth to that of the DP **(Figure 4P)**. While morphogen-driven growth of individual tissues can be developmentally regulated, tissue- or layer-intrinsic morphogen signaling cannot account for adaptive, synchronized growth of two or more tissues or tissue layers, because adaptive growth coordination fundamentally requires communication of growth information between tissues. A shift from morphogen-dependent to mechanically-dependent growth in one tissue layer might be a general mechanism that facilitates coordinated growth. Such coordination of growth between different tissues in an organ might be a key function of the Hippo pathway under normal physiological conditions.

## Materials and Methods

### RESOURCE AVAILABILITY

#### Lead contact

Further information and requests for resources and reagents should be directed to and will be fulfilled by the lead contact, Iswar K. Hariharan.

#### Materials availability

This study did not generate new unique reagents. However, the *GMR43F06-GAL4* fly line is no longer available from the Bloomington *Drosophila* Stock Center and is available upon request.

#### Data and code availability

Microscopy data reported in this paper will be shared by the lead contact upon request. Code used for analyses in R is available through public repositories as indicated in the Methods section of this manuscript.

### EXPERIMENTAL MODEL AND SUBJECT DETAILS

#### Drosophila maintenance

*Drosophila* were maintained at room temperature (approx. 22°C) on standard molasses food supplemented with yeast paste. Timed egg lays were performed at 25°C on grape juice-agar plates supplemented with yeast paste, after which first-instar larvae were transferred to molasses food with yeast paste at a density of 50-60 larvae per vial. For experiments involving precise developmental timing, larvae were raised in a 25°C incubator unless otherwise specified. “Wild type” denotes Oregon-R (OreR) flies unless otherwise stated.

### METHOD DETAILS

#### Induction of clones

To generate *tkv^4/4^* clones, *tkv^4^ FRT40A / CyO* males were crossed with *hsFlp; GFP FRT40A/CyO* virgin females. From these crosses, larvae were heat shocked at 37°C at 24±2h AEL for 1h, then raised at 25°C until dissection in late L3.

To generate *ban^Δ1/Δ1^* clones, *Sp/CyO; ban^Δ1^ FRT80B/TM* males were crossed with *hsFlp; GFP FRT80B/TM3* virgin females. From these crosses, larvae were heat shocked at 37°C at 60±12h AEL for 1h, then raised at 25°C until dissection in late L3.

#### Timing of temperature-sensitive experiments

For all timed experiments, egg lays were performed for 4 h at 25°C.

For the *stripe>smo-RNAi* “RNAi always on” condition, eggs were moved to 30°C immediately following the egg lay and resulting larvae were raised entirely at 30°C until dissection in the late third instar. For the *stripe^ts^>smo-RNAi* “RNAi on from mid-embryogenesis” condition, eggs were incubated at 18°C from immediately after the egg lay until 24±2h AEL (approximately equivalent to ∼12h AEL at 25°C), at which time they were moved to 30°C until dissection in the late third instar. For the *stripe^ts^>smo-RNAi* “RNAi on from ∼hatching” condition, eggs were incubated at 18°C from immediately after the egg lay until 48±2h AEL (approximately equivalent to ∼24h AEL at 25°C), at which time they were moved to 30°C until dissection in the late third instar.

For the *hh^ts^/hh^AC^* and *hh/hh^+^*experiments shown in **S2C-D**, eggs and resulting larvae were incubated at 18°C from immediately after the egg lay until 140±2h AEL, at which time they were moved to 30°C until dissection in the third larval instar at 168±2h AEL. For the *hh^ts^/hh^AC^*18°C control shown in **S2F**, eggs and resulting larvae were incubated at 18°C until dissection in the late third instar.

For the *PE^ts^>GFP, yki-RNAi* experiment shown in **3F**, unstaged eggs and resulting larvae were raised at 18°C for ∼4 days before being transferred to 30°C until dissection in the late third instar.

#### Immunohistochemistry

Imaginal discs were dissected in phosphate buffered saline (PBS) and fixed in PBS with 4% paraformaldehyde at 22°C for 20 min. Discs were washed 3X 5 min wash with PBS containing 0.1% Triton X-100 (PBST). Discs were incubated for 30 min in PBST containing 10% normal goat serum, except for discs stained for Hth protein, which were incubated in PBST containing 10% normal donkey serum. Discs were then incubated in PBST containing 10% normal goat serum (or donkey serum for Hth stains) containing primary antibodies overnight at 4°C. Discs were washed 3X 10 min in PBST. Discs were then incubated in PBST containing 10% normal goat serum (or donkey serum for Hth stains) and the appropriate goat (or donkey) Alexa Fluor-conjugated secondary antibody (1:500; Invitrogen) for 2 h. Discs were incubated in PBT with DAPI (1:1000) for 20 minutes, washed quickly 3X in PBS, and mounted in Slowfade Diamond Antifade Mountant (Invitrogen) on slides with Scotch Magic Tape spacers.

Primary antibodies used: mouse anti-Eya (1:100; DSHB), rabbit anti-phospho-histone-H3 (1:500; Millipore), rabbit anti-phospho-Smad3 (1:500; abcam), rat anti-Ci (1:25; DSHB), chick anti-GFP (1:500, abcam), rabbit anti-Dcp-1 (1:250; Cell Signaling), goat anti-Hth (1:50; Santa Cruz Biotechnology), rat anti-E-cad (1:100; DSHB), and mouse anti-Mmp1 (1:100 each of 3 antibodies; DSHB). Further antibody details are provided in the Key Resources Table of this manuscript.

#### EdU staining

EdU labeling was performed using an Invitrogen Click-iT EdU 647 Imaging Kit. Half carcasses were dissected and inverted in Schneider’s *Drosophila* Medium, then incubated with rocking in 10 μM EdU in Schneider’s Drosophila Medium at 22°C for 60 min. Tissues were fixed in 4% paraformaldehyde in PBS for 20 min, then quickly washed twice with PBST and permeabilized in PBS with 0.3% Triton for 20 min. Tissues were quickly washed twice in PBS, then incubated in Click-iT reaction cocktail (prepared in accordance with manufacturer instructions) at 22°C for 30 min, protected from light. Tissues were washed twice with PBS. Subsequent antibody staining proceeded as described above, starting with the 30 min incubation in PBST containing 10% normal goat serum.

#### Microscopy and image processing

Adult wings were mounted in Gary’s Magic Mountant. Images of adult flies and adult wings were obtained on a Keyence VHX-5000 digital microscope. Adult wing measurements were performed on female wings exclusively, due to size differences between male and female *Drosophila*.

Imaginal discs were imaged with a Zeiss Axio Imager.M2 fluorescence microscope equipped with a Zeiss Apotome.2 Structured Illumination device to perform optical sectioning. Apotome images were deconvolved in Zen 2.3 Pro.

All images were processed in FIJI (Schindelin et al., 2012). For wing discs that were larger than the image frame, multiple images were combined using the Stitching plugin in FIJI (Preibisch et al., 2009).

### QUANTIFICATION AND STATISTICAL ANALYSIS

All statistical analyses were performed in R using open-source software. Unpaired t-tests were performed in R. Exact values of n can be found in the figure legends. A cutoff of p<0.05 was used to define statistical significance.

Scatter plots were generated in Excel. Bar charts were generated using the ggplot2 package in RStudio (Wickham, 2016). Bar charts represent average values and error bars represent the standard deviation.

#### Cell area and fluorescence intensity measurements

Cell area in the most squamous region of the wing disc was calculated by selecting a 50 μm x 50 μm square in the squamous PE and dividing the area of the square by the number of nuclei it contained.

Fluorescence intensity was measured in FIJI within the plane of the PE (or the DP, when appropriate), and represents the average intensity over the entire region of the disc unless otherwise stated.

#### Single cell data analysis

Single-cell RNA sequence data (Everetts et al., 2021) represents harmonized data from 120h AEL and 96h AEL wing discs. Data was processed using Seurat (Hao et al., 2021), following the standard workflow described in the Seurat guided clustering tutorial (https://satijalab.org/seurat/articles/pbmc3k_tutorial.html).

For quality control, cells with fewer than 200 or greater than 2500 features were removed from the data; it was then log normalized and reduced to 3000 variable features, and PCA was performed. Harmony (Korsunsky et al., 2019) was used to minimize batch effects from the different time points.

After clustering was performed in Seurat, we used expression of *FasIII* (a disc epithelium marker) and lack of expression of *twi* (a myoblast marker) to identify clusters corresponding to disc epithelium identity. We restricted further gene expression analyses to the population identified as disc epithelial cells. We identified the cell cluster corresponding to PE identity by high, specific expression of the PE marker genes *Ubx, eya,* and *so*. Gene expression dot plots were generated in RStudio using the Seurat DotPlot function.

### KEY RESOURCES TABLE

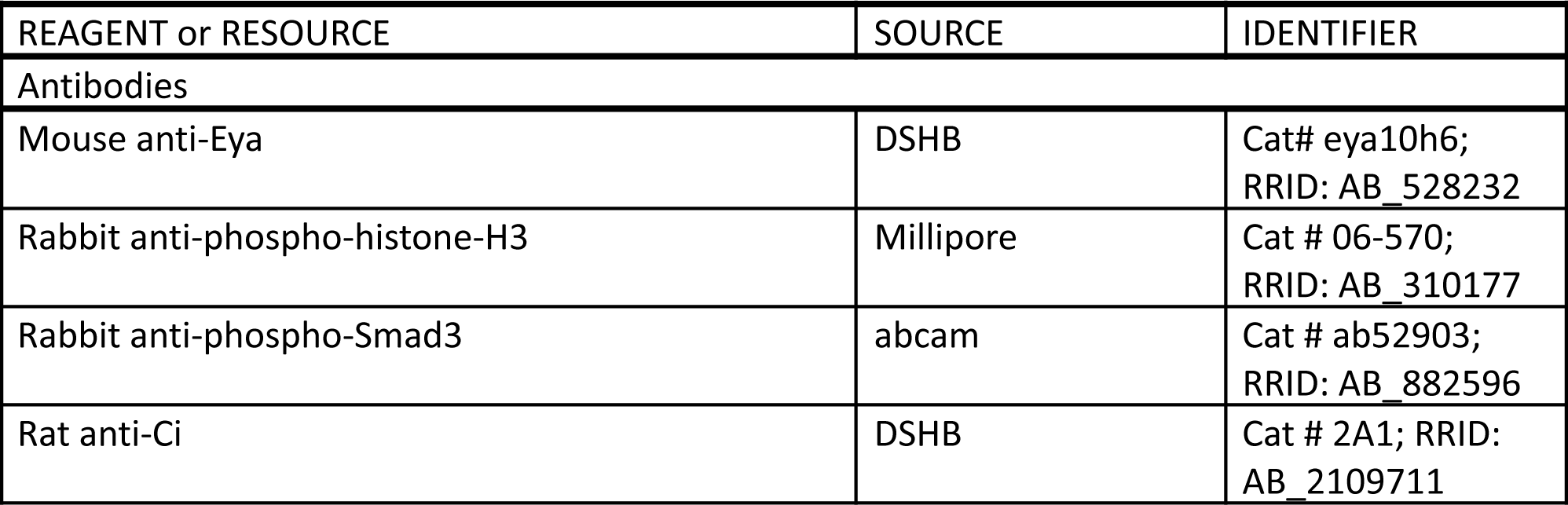

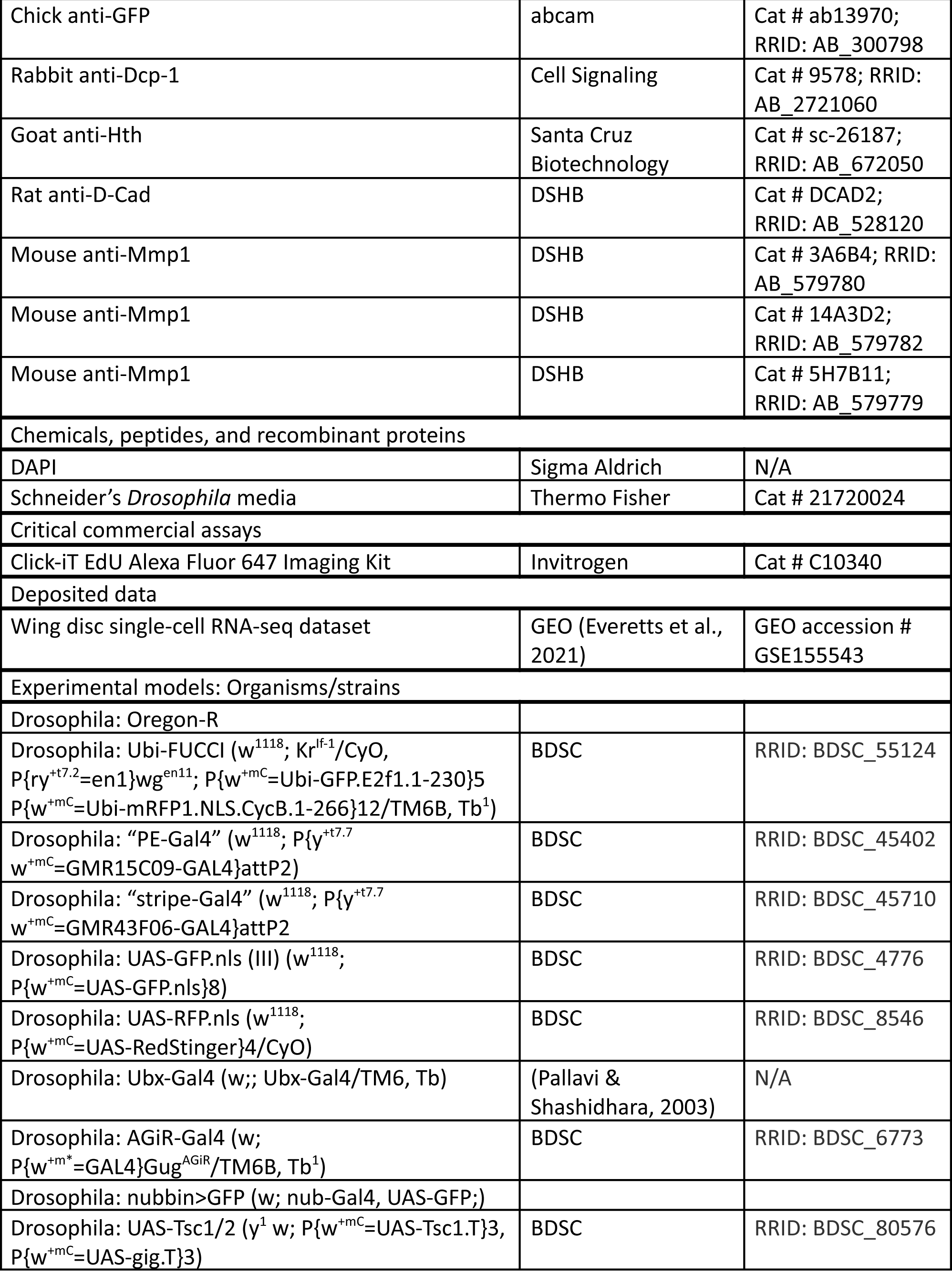

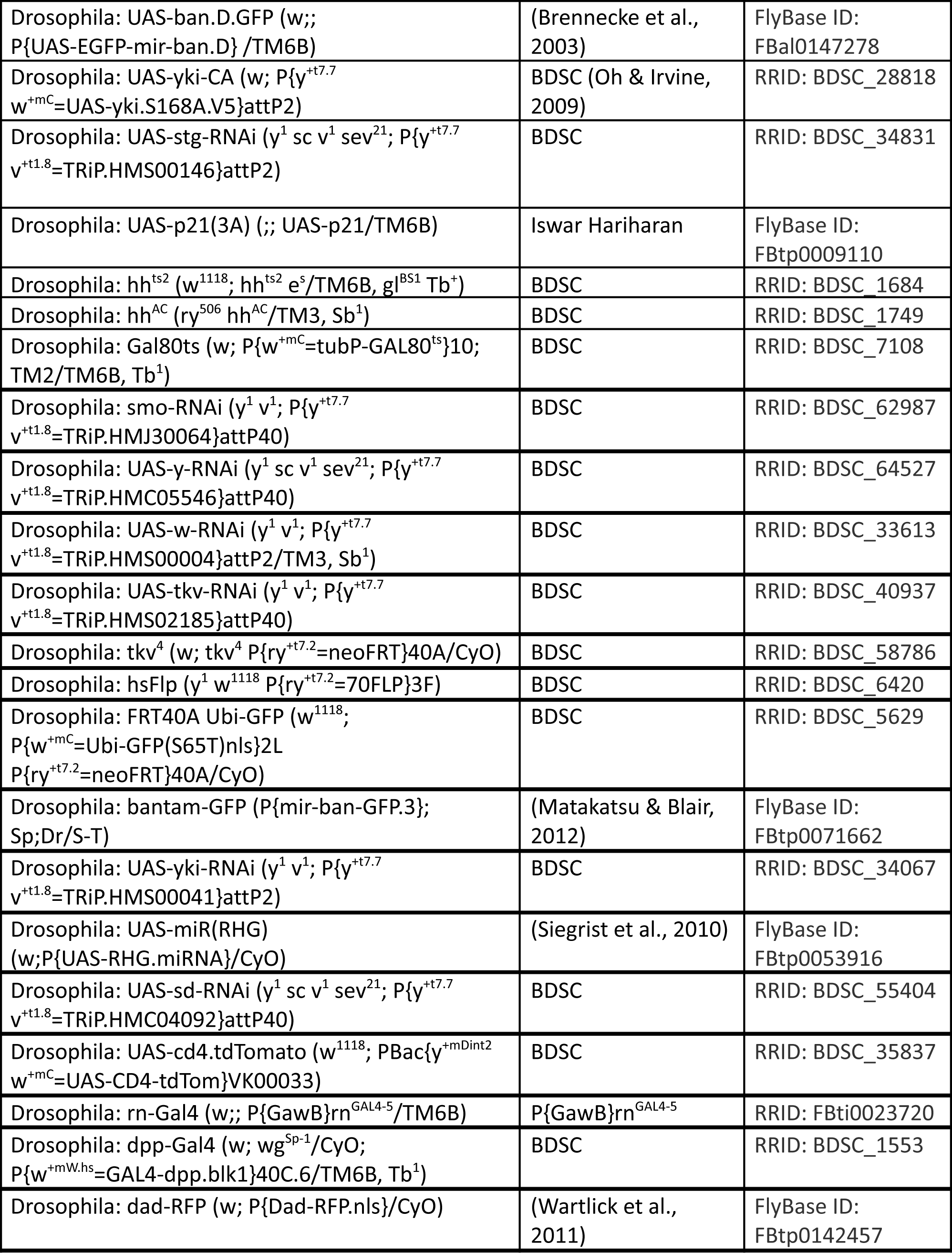

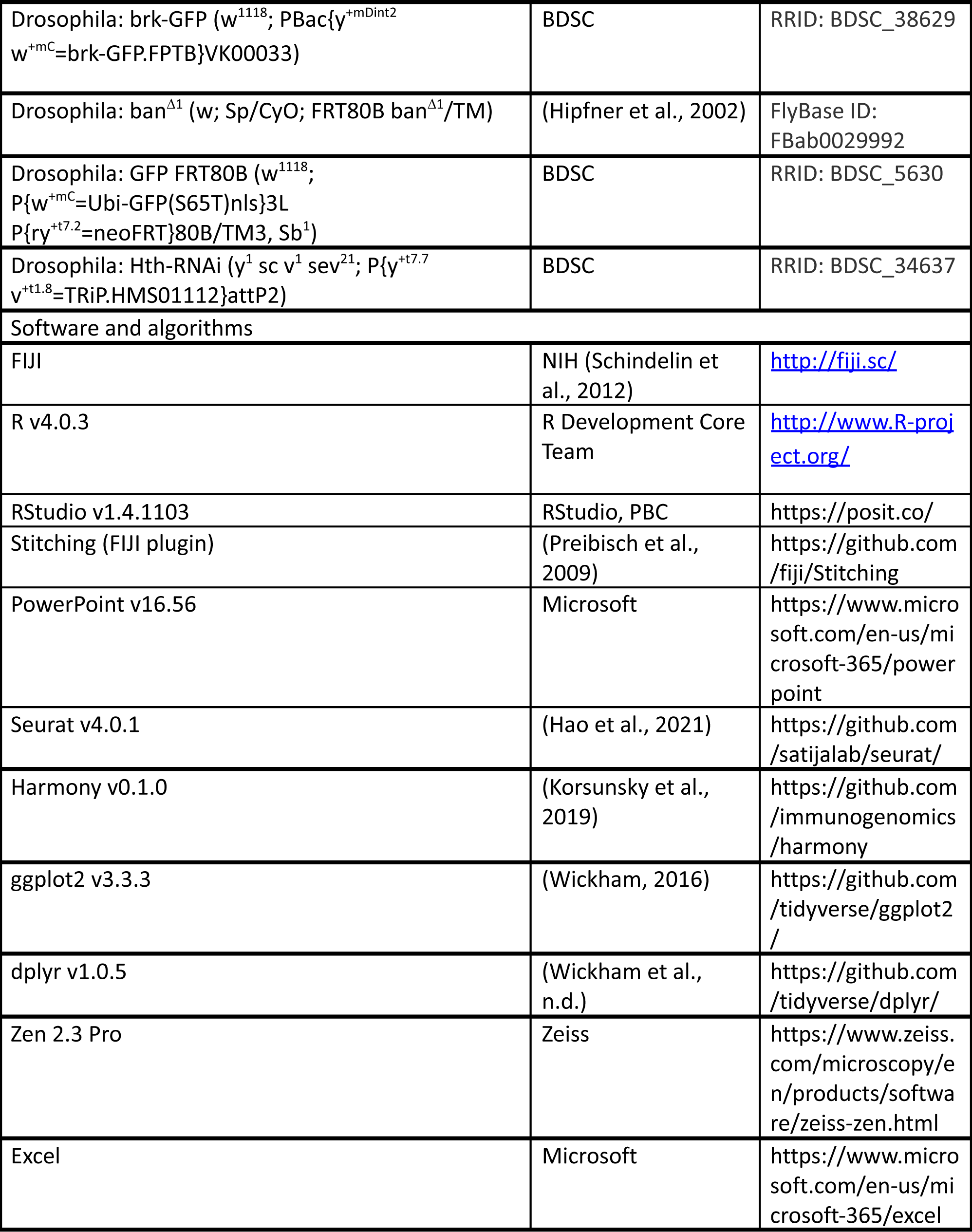

## Acknowledgments

The authors would like to thank Stephen Cohen, Ajay Srivastava, Nicolas Tapon, and Richard Mann for generously sharing stocks, current and former members of the Hariharan lab for invaluable feedback, Noah Whiteman, Craig Miller, and David Bilder for scientific advice, Melanie Worley for comments on the manuscript, and the Bloomington Stock Center, DRSC/TRiP Functional Genomics Resources, and Developmental Studies Hybridoma Bank for stocks and reagents.

## Supplemental Figures

**Figure S1.**
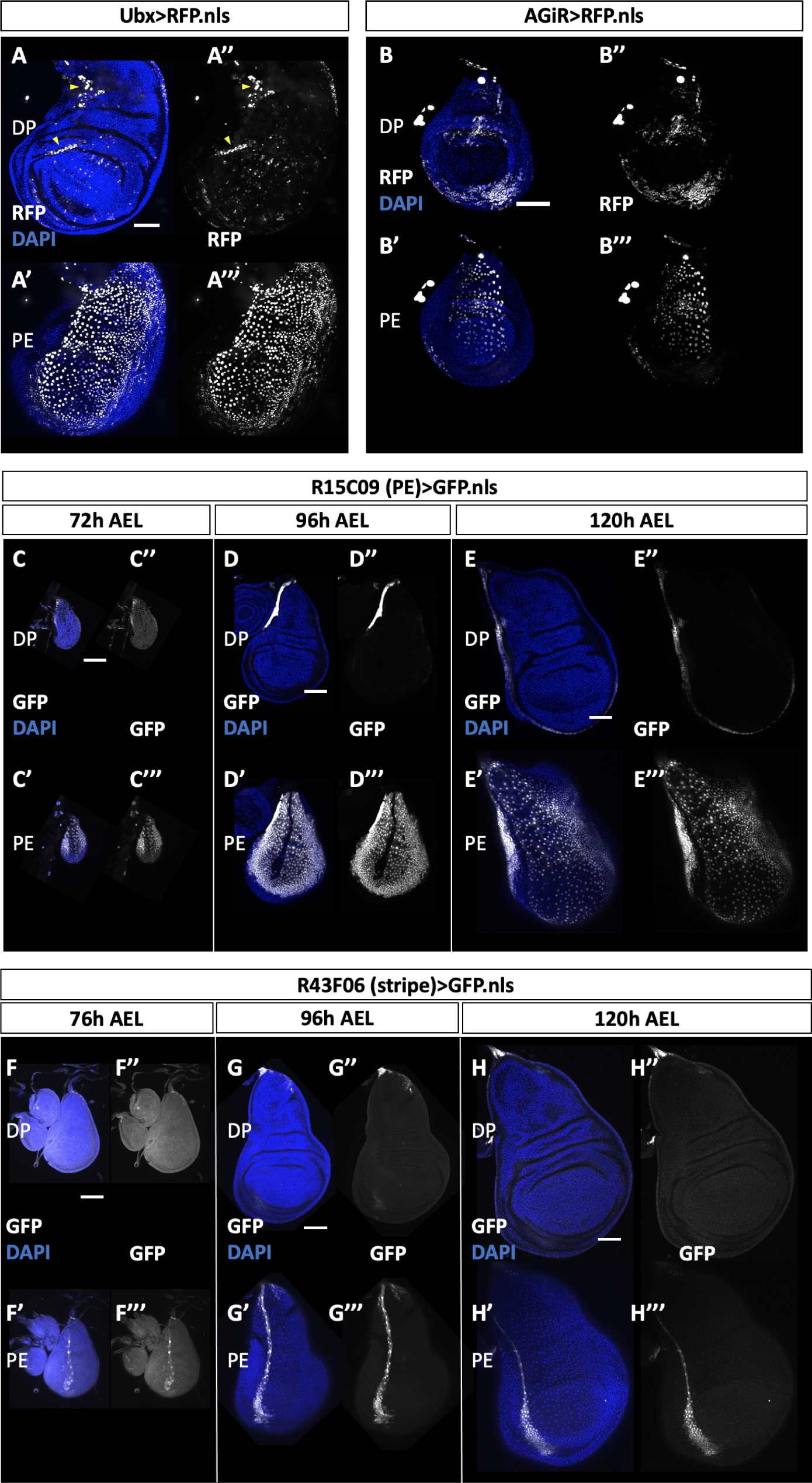
Characterization of PE-specific drivers. **A)** Expression of the *Ubx-Gal4* driver, as labeled by UAS-RFP (white), includes squamous PE cells as well as a “salt-and-pepper” pattern of cells within the DP. Arrowheads mark regions where folding of the disc has pushed regions of the PE into the plane of the DP. **B)** Expression of the *AGiR-Gal4* driver, as labeled by UAS-RFP (white), includes most squamous PE cells in addition to a large population of cells within the hinge region of the DP. **C)** Expression of the *R15C09-Gal4* driver (also referred to as *PE-Gal4*) is largely specific to the PE throughout L3. Image is of a disc 72h AEL. Note that GFP expression (white), marking driver activity, is exclusive to the PE with the exception of a small region at the extreme anterior margin of the DP, contiguous with PE expression. **D)** A mid-L3 wing disc (96h AEL) showing PE-specific expression of *PE-Gal4*. **E)** A late-L3 wing disc (120h AEL) showing PE-specific expression of *PE-Gal4*. **F)** Expression of the *R43F06-Gal4* driver (also referred to as *stripe-Gal4*), as labeled by UAS-GFP (white), is largely specific to the anterior-posterior compartment boundary of the PE throughout L3. A small region of cells in the “stem” of the disc also express the driver. An early-L3 wing disc (72h AEL) showing *stripe-Gal4* expression. **G)** A mid-L3 wing disc (96h AEL) showing PE-specific expression of *stripe-Gal4*. **H)** A late-L3 wing disc (120h AEL) showing PE-specific expression of *stripe-Gal4*. All scale bars are 100µm.

**Figure S2.**
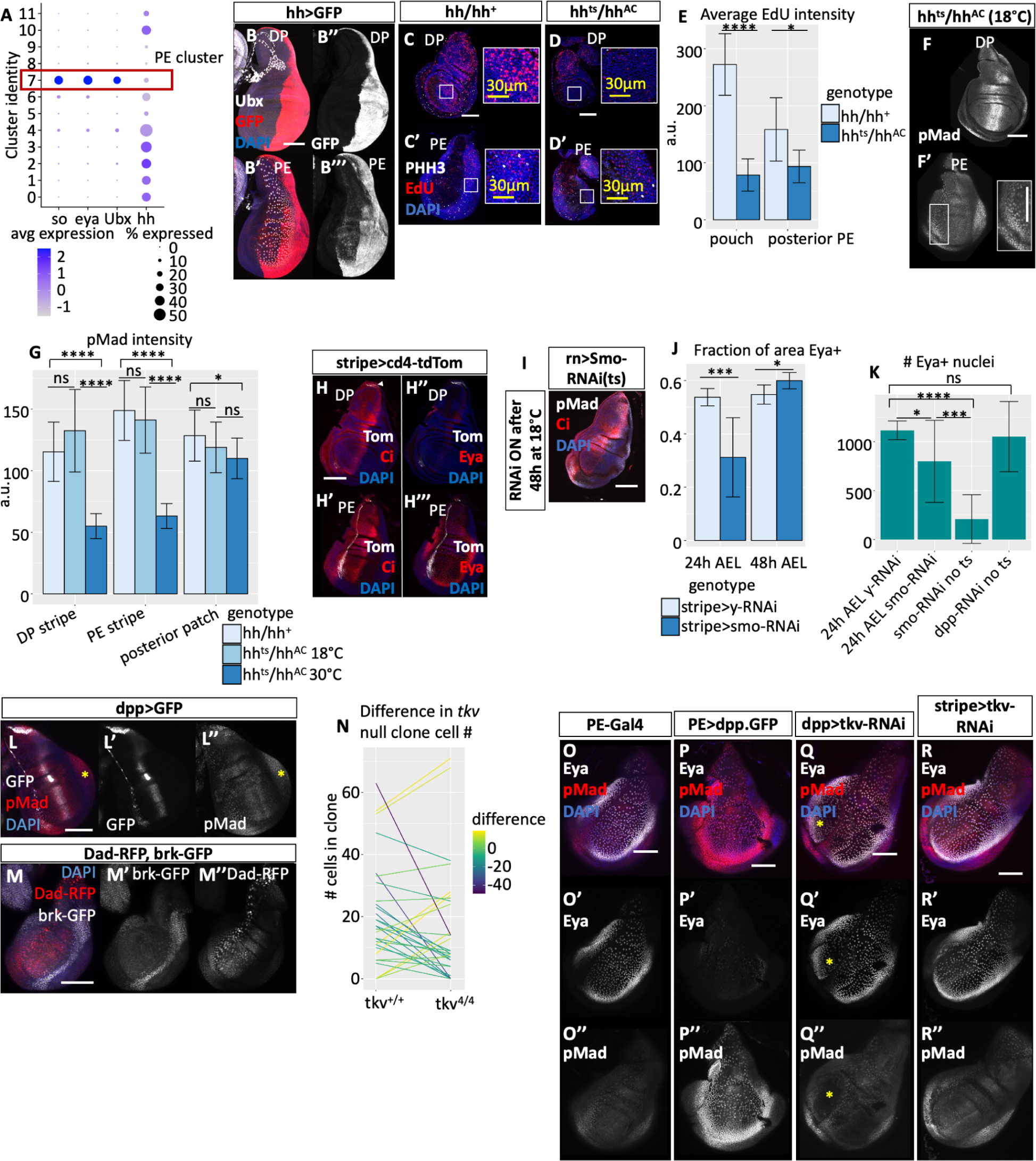
Supplemental data on Hh and Dpp. **A)** *hh* expression is low, but present, within the wing disc PE. Within a wing disc scRNAseq dataset(Everetts et al., 2021), we identified disc epithelial cells by expression of *FasIII* and lack of expression of *twi,* and further identified PE cells by expression of the PE marker genes *so, eya,* and *Ubx* (details in Materials and Methods). **B)** *hh-Gal4* is expressed in the posterior PE. anti-GFP (red in left panels, white in right panels) was used to detect faint GFP expression. The strong staining is bleedthrough from the DP. **C)** Global loss of Hh function, caused by exposing larvae with only a temperature-sensitive *hh* allele to the nonpermissive temperature, reduces proliferation in both layers of the wing disc. Proliferation is observed by EdU+ incorporation (red) and expression of phospho-histone H3 (white). Control discs, which have one functional *hh* allele, show robust proliferation at the nonpermissive temperature (30°C). **D)** *hh^ts^/hh^AC^* wing discs at the nonpermissive temperature show reduced proliferation, as observed by the reduced number of EdU+ puncta (red) and pHH3+ cells (white). **E)** Quantification of EdU intensity from the conditions shown in **(C-D)**. Average EdU intensity was measured from a representative 50×50 μm square region within the plane of the DP or PE, as appropriate. n = 8 discs (*hh/hh^+^*), 5 discs (*hh^ts^/hh^AC^*). Unpaired t-test of average EdU intensity between *hh/hh^+^* and *hh^ts^/hh^AC^* ****p<0.00001 in the pouch of the DP, *p<0.05 in the posterior PE. **F)** At the permissive temperature (18C), larvae that express only temperature-sensitive *hh* still have a stripe of Dpp signaling (visualized by pMad, white) at the AP compartment boundary of both layers. Discs at the non-permissive temperature are shown in Figure 3C. **G)** Quantification of pMad intensity at the AP boundary of the DP (“DP stripe”), the AP boundary of the PE (“PE stripe”), and the posterior patch of Dpp signaling in the PE upon global loss of Hh function. At the nonpermissive temperature (30°C), the *hh^ts^*mutant (dark blue) greatly reduces Dpp signaling at the AP boundary in both layers, compared to both a temperature-matched hh heterozygote (*hh/hh^+^*, pale blue) and to the temperature-sensitive mutant at the permissive temperature (18°C). The posterior patch of Dpp signaling is minimally affected by Hh loss (Foronda 2009). n = 12 discs (*hh/hh^+^*), 12 discs (*hh^ts^/hh^AC^*). Unpaired t-tests of pMad intensity between *hh/hh^+^* and 18°C control are ns (not significant) for all regions. Unpaired t-tests of pMad intensity between *hh^ts^/hh^AC^*and 18°C control or between *hh^ts^/hh^AC^* and *hh/hh^+^*are ****p<0.00001 in both the DP stripe and the PE stripe. Unpaired t-test of pMad intensity in the posterior patch between *hh^ts^/hh^AC^*and 18°C control ns (not significant). Unpaired t-test of pMad intensity in the posterior patch between *hh^ts^/hh^AC^* and *hh/hh^+^*\*p<0.05. **H)** The *stripe-Gal4* driver has minimal expression in the DP in L3 (seen by cd4-tdTomato expression, white, arrowhead). In the PE, *stripe-Gal4* expresses along the AP compartment boundary, mostly within the anterior compartment (seen by Ci expression, red in left panels) but including a few cells just posterior to the compartment boundary. *stripe-Gal4* expression is approximately inverse to Eya expression (red in right panels). **I)** Loss of Hh signaling within the pouch of the DP, caused by knocking down *smo* using *rotund-Gal4 (rn-Gal4)*, results in dramatic tissue undergrowth. Knockdown was initiated at 48h AEL at 18°C. **J)** *smo* knockdown causes PE undergrowth, as measured by the fraction of the total disc area that is Eya+, when knockdown begins at 24h AEL, but not when knockdown begins at 48h AEL. *y-RNAi* (control) and *smo-RNAi* were expressed with *stripe-Gal4.* n = 10 discs (24h AEL *y-RNAi*), 10 discs (24h AEL *smo-RNAi*), 16 discs (48h AEL *y-RNAi*), 8 discs (48h AEL *smo-RNAi*). Unpaired t-test of Eya+ fraction between 24h AEL *y-RNAi* and 24h AEL *smo-RNAi* ***p<0.001; unpaired t-test of Eya+ fraction between 48h AEL *y-RNAi* and 48h AEL *smo-RNAi* *p<0.05. **K)** *smo* knockdown reduces the number of Eya+ cells in the PE, especially when knockdown begins very early. *dpp-RNAi* does not change the number of Eya+ cells in the PE. “24h AEL” indicates that knockdown was initiated after 24h of development at 18°C; “no ts” indicates that larvae do not express Gal80^ts^. All knockdowns were expressed with *stripe-Gal4.* n = 5 discs (24h AEL *y-RNAi*), 11 discs (24h AEL *smo-RNAi*), 11 discs (*smo-RNAi* no ts), 7 discs (*dpp-RNAi* no ts). Unpaired t-test of the number of Eya+ nuclei between 24h AEL *y-RNAi* and 24h AEL *smo-RNAi* *p<0.05; unpaired t-test of the number of Eya+ nuclei between 24h AEL *y-RNAi* and *smo-RNAi* no ts ****p<0.00001; unpaired t-test of the number of Eya+ nuclei between 24h AEL *smo-RNAi* and *smo-RNAi* no ts ***p<0.001; unpaired t-test of the number of Eya+ nuclei between 24h AEL *y-RNAi* and *dpp-RNAi* ns (not significant, p = 0.66). **L)** *dpp-Gal4* is expressed in the PE in an analogous pattern to that of the DP. Note stripe of *dpp>GFP* expression (white in **L** and **L’**), surrounded by a broader region of Dpp signaling activity (pMad, red in **L**, white in **L’’**). Asterisk marks posterior Dpp signaling that is Hh-independent and that is not captured by the expression of *dpp-Gal4*. **M)** The canonical Dpp target genes *Dad* (expression promoted by Dpp signaling) and *brk* (repressed by Dpp signaling) have patterned expression in the PE, consistent with a role for PE Dpp signaling in patterning the PE. Expression was observed by *dad-RFP* (red in **M**, white in **M’**) and *brk-GFP* (white in **M** and **M’’**). **N)** *tkv* null clones (*tkv^4^/tkv^4^*) usually have fewer cells than their wild-type twin spots. n = 31 clone pairs across 5 discs. **O)** Eya expression in the wild-type PE (white in **O** and **O’**) is approximately inverse to PE Dpp signaling activity as measured by pMad (red in **O**, white in **O’’**). **P)** Dpp overexpression in the PE using *PE>dpp.GFP* causes increased PE pMad (red in **P,** white in **P’’**) and dramatically decreased Eya expression (white in **P** and **P’**). **Q)** *dpp>tkv-RNAi* reduces Dpp signaling at the AP boundary of the PE (pMad, red in **Q,** white in **Q’’**; note gap in pMad staining at yellow asterisk), and causes ectopic Eya expression (white in **Q** and **Q’**; note additional expression at yellow asterisk). **R)** Reducing Dpp signaling at the AP boundary of the PE using *stripe>tkv-RNAi* similarly causes additional Eya expression (white in **R** and **R’**; note narrower gap of Eya expression compared to **O’**). All scale bars are 100µm unless otherwise stated.

